# Multiplex imaging of localized prostate tumors reveals changes in mast cell type composition and spatial organization of AR-positive cells in the tumor microenvironment

**DOI:** 10.1101/2023.08.18.553854

**Authors:** Cigdem Ak, Zeynep Sayar, Guillaume Thibault, Erik A. Burlingame, Jennifer Eng, Alex Chitsazan, Andrew C. Adey, Christopher Boniface, Paul T. Spellman, George V. Thomas, Ryan P. Kopp, Emek Demir, Young Hwan Chang, Vasilis Stavrinides, Sebnem Ece Eksi

## Abstract

Mapping spatial interactions of cancer, immune and stromal cells present novel opportunities for patient stratification and for advancing immunotherapy. While single-cell studies revealed significant molecular heterogeneity in prostate tumors, there is currently no understanding of how immune cell heterogeneity impacts spatial coordination between tumor and stromal cells in localized tumors. Here, we used cyclic immunofluorescent imaging on whole-tissue sections to uncover novel spatial associations between cancer and stromal cells in low- and high-grade prostate tumors and tumor-adjacent normal tissues. Our results provide a spatial map of 699,461 single-cells that show epigenetic and molecular differences in distinct clinical grades. We report unique populations of mast cells that differentially express CD44, CD90 and Granzyme B (GZMB) and demonstrate GZMB+ mast cells are spatially associated with M2 macrophages in prostate tumors. Finally, we uncover recurrent neighborhoods that are primarily driven by androgen receptor positive (AR+) stromal cells and identify transcriptional networks active in AR+ prostate stroma.

## MAIN

Prostate tumors are composed of epithelial cells that exhibit molecular and cellular differences and interact with a complex ecosystem of stromal cells: fibroblasts, immune cells, mesenchymal stem cells, smooth muscle, blood vessels and innervating neurons [1]. The complexity and heterogeneity of disease is indicated by pathological and single-cell omic evidence [2-4]. It is essential to simultaneously dissect this cellular heterogeneity and generate a spatial map between cell types and states to better stratify patients into treatment groups and to identify more effective immunotherapy strategies. Not surprisingly, these cell states, types and bidirectional cellular interactions may not be solely recovered from Hematoxylin and Eosin (H&E) staining, even with advanced machine learning algorithms. Single-cell -omic approaches can identify cell types and states, but not the spatially coordinated behavior of heterogenous cells. Immunohistochemistry is gaining widespread adoption in clinical practice to mitigate this problem, but it is often deployed for a few markers in a non-systematic, non-standardized manner.

Recent advancements in multiplex proteomic profiling and image analysis facilitated the profiling of >30 proteins from thousands of single-cells on single tissue-sections, opening a new path in predicting disease subtypes based on immune microenvironments, linked to progression [5, 6]. In breast cancer, structured immune microenvironments linked to survival were identified [7-9]. In colorectal cancer, distinct cellular communities, which can distinguish molecular subtypes and aid in the risk-stratification of aggressive forms of cancer were revealed [10]. These studies show the power of mapping spatial associations in the tumor microenvironment (TME) for predicting patient trajectory and revealing novel targets for therapeutic interventions [4].

In the prostate TME, dynamic changes in the physical microenvironment of prostate tumors, such as hypoxia, wound healing and chronic inflammation are postulated to contribute to cancer initiation and progression [11-13]. These processes require coordination between immune and non-immune stromal cells. However, our understanding of non-immune stromal cell subtypes is limited, especially around AR expression in prostate TME. A few studies report loss of stromal AR expression that is strongly correlated with the worst patient outcomes [14, 15]. However, a better delineation of grade-specific AR+ stromal cell-types and of pathways involved in AR regulation in stromal cells is needed [16].

Here, we developed an extensive antibody library to probe changes in the prostate TME and identified 27 epithelial, immune and non-immune stromal cell phenotypes that exhibit prostate cancer grade-specific differences in spatial composition (Figure 1a). We used cyclic immunofluorescent (cycIF) imaging to spatially map 699,461 cells from 13 patients and three clinical-grades (Figure 1a). Further, we analyzed copy number aberrations (CNA) for 10 patients to investigate changes in the spatial coordination of immune and non-immune stromal cells. We show AR+ stroma driven neighborhood organization associated with clinical grade and defined transcriptional networks active in AR+ stroma using single-cell ATAC sequencing (sci-ATAC-seq) from the same group of patients [17]. Our study provides a comprehensive spatial and regulatory map of the prostate TME for patients with localized prostate tumors and highlights the importance of identifying disease subtypes for improving patient outcomes.

**Figure 1:**
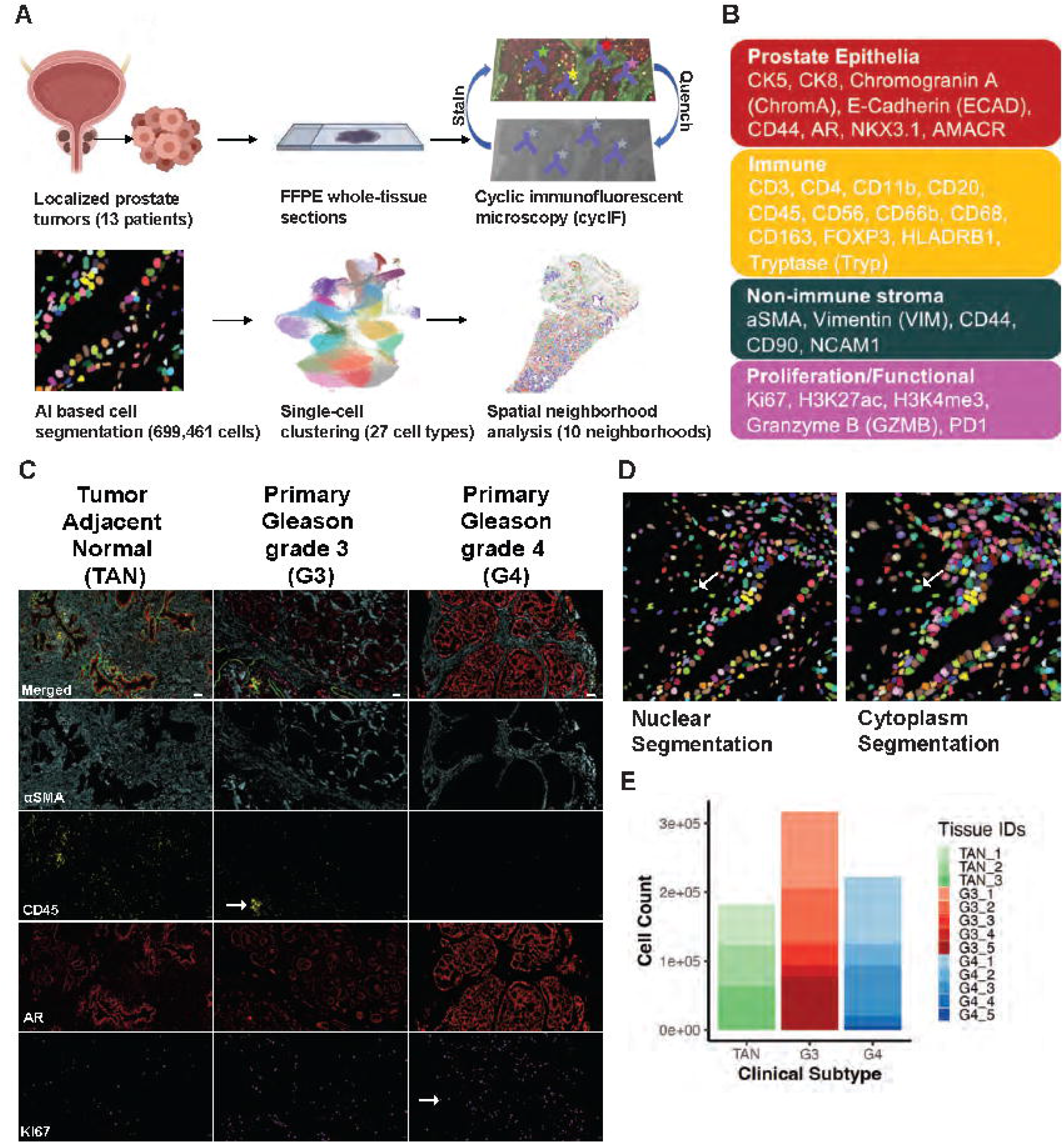
Subcellular profiles of 28 proteins are captured from 699,481 single-cells via cyclic immunofluorescent imaging (cycIF). (A) Experimental and computational pipeline. FFPE sections were collected from three tumor adjacent normal (TAN), five primary Gleason grade 3 (G3) and five primary Gleason grade 4 (G4) treatment-naïve, localized prostate tumors. Tissue-sections were stained, imaged, quenched and imaged for 10 rounds. Cell segmentation detected 699,461 cells in total that were clustered into 27 cell subtypes and 10 cell neighborhoods. (B) Antibody library used in the study distinguishes between epithelial (red), immune (yellow) and non-immune stromal cells (teal) and includes several functional status markers (pink) for each cell type. (C) Example cycIF images from TAN, G3 and G4 tissue-sections show αSMA (blue), CD45 (yellow), AR (red) and Ki67 (magenta) markers. Arrows point out to decreasing immune populations with grade (CD45) and increasing population of proliferating cells (Ki67). (D) Two-step segmentation step (nuclei and cell segmentation) that determines the boundaries of each cell. Arrows point out to the defined boundaries of a stromal cell. (E) Numbers of cells were segmented from each clinical grade TAN, G3 and G4.

## RESULTS

### Developing a prostate-cancer specific antibody library to probe immune and non-immune stromal cell type changes

To study systemic changes in the prostate TME, we developed an extensive antibody library to capture prostate cancer cells in relation to immune and non-immune stromal cell subtypes. We tested over 150 antibodies and validated more than 100 antibodies following a three-step verification for each antibody (Figure S1, Supplementary Files 1 and 2). Our antibody library is specifically geared toward capturing changes in prostate cancer biology.

We selected 28 targets from this panel for cycIF, which allowed for a thorough classification of specific cell types in the prostate tissue: epithelial (CK5, CK8, E-Cadherin (ECAD), Chromogranin A (ChromA) and CD44), stromal (αSMA, Vimentin (VIM), CD44, CD90, NCAM) and immune cells (CD3, CD4, CD11b, CD20, CD45, CD56, CD66b, CD68, CD163, FOXP3, HLADRB1, Tryptase (Tryp)). We also included molecular and epigenetic markers (AR, NKX 3.1, AMACR, H3K27ac, H3K4me3), immune cell functional status markers (Granzyme B (GZMB), PD1) and a proliferation marker to capture specific cell states associated with patient outcome (Ki67) (Figure 1b, Figure S2).

### Profiling spatial localization of 28 proteins from 699,461 cells isolated from tumors spanning three clinical grades

FFPE whole-tissue sections from 13 patients with localized primary Gleason Pattern 3 (G3) or Gleason Pattern 4 (G4) prostate tumors were obtained for cycIF analysis (Figure 1c). Whole-slide images varied between 2 to 26mm^2^ in surface area. We did not generate tumor microarrays (TMAs) to avoid any sampling bias [18]. Three tissue-sections where >95% of the section present were histopathologically normal glands or stroma and less than <5% of the section was Gleason pattern 3 were assigned as ‘tumor adjacent normal’ (TAN). Five tissue-sections where >75% of the section consisted of Gleason pattern 3 tumors were assigned as ‘Gleason pattern 3’ (G3) and five tissue-sections where >75% of the section consisted of Gleason pattern 4 tumors were assigned as ‘Gleason pattern 4’ (G4) (Figure S3). The Cancer of the Prostate Risk Assessment Scores (CAPRA-S) scores for all patients were calculated based on clinical patient data [19]. We identified 5 patients with low, 6 patients with intermediate and 2 patients with high CAPRA-S risk.

We analyzed the co-expression of 28 proteins across 13 tissues using cycIF. Single-cell segmentation via CellPose identified 699,461 cells, with similar numbers of cells from each clinical grading (G3, G4 and TAN) (Figure 1d-e) [20].

### The density of epithelial, immune and non-immune stromal cell types indicates changes in the cellular organization of tissues associated with clinical-grade

Our previous work uncovered a loss of epigenomic heterogeneity in epithelial prostate cancer cells [17]. To further investigate cell type heterogeneity, we performed a two-step clustering analysis for an in-depth classification of epithelial, immune and non-immune stromal cells in prostate tumors. First, we clustered all 699,461 single-cells based on the expression of every marker and detected 49 clusters (Figure 2a) [21]. We used UMAP to visualize single-cells based on their 28-plex protein expression data, colored by Louvain cluster identities (IDs), patient IDs matched with primary Gleason pattern and broad cell type categories (Figure 2a) [22]. Single-cell clusters on the UMAP space showed patient-specific epithelial cell clusters (Figure 2a). In contrast, immune and non-immune stromal cells across all patients formed a single, mixed cluster separate from all epithelial cells (Figure 2a).

**Figure 2:**
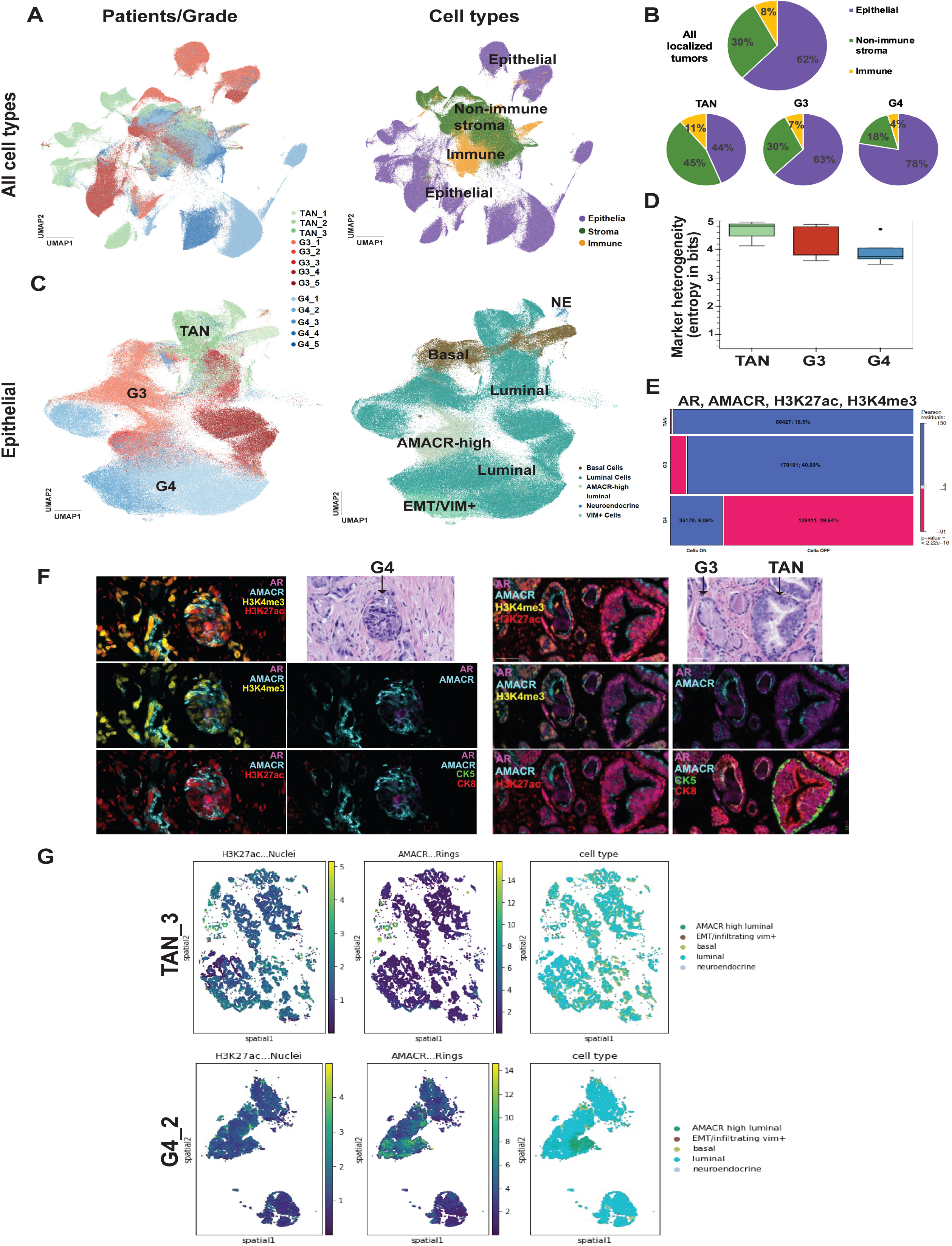
Combined expression of four markers— AR, AMACR, H3K27ac and H3K4me3— distinguish G4 tumors. (A) Louvain clustering of 699,481 cells identified clinical-grade specific (left) and cell-type specific (right) clusters. (B) Fraction of epithelial cells (green) increased and stromal cells (magenta and red) decreased in G4 tumors. (C) Louvain clustering of epithelial cells shows cells from G3 tumors form transitionary clusters between TAN and G4 samples and VIM+ epithelial cells cluster away from basal cells. (D) Shannon entropy results measuring the distribution of cells that express epithelial markers across clinical subtypes which shows a decrease in the marker heterogeneity with increasing tumor grade with a not significant p-value. (E) Combined expression of four markers—AR, AMACR, H3K27ac and H3K4me3— distinguish between G4 samples from TAN and G3 samples (chi-square test with p<2.2e-16). This mosaic plot with colored cases shows where the observed frequencies deviates from the expected frequencies. The pink cases means that the observed frequencies are smaller than the expected frequencies, whereas the blue cases means that the observed frequencies are larger than the expected frequencies. The area of the box is proportional to the difference in observed and expected frequencies. Moreover, p-value of the Chi-square test is also displayed at the bottom right. (F) A G4 gland detected by an H&E show expression of AR, AMACR, H3K27ac and H3K4me3 but does not express CK8 (luminal) or CK5 (basal) markers (left) A G3 and normal gland detected by an H&E show CK8 and CK5 expression. (G) Plot showing distribution of H3K27ac and AMACR in TAN (top) vs. G4 (bottom) samples.

To delineate cellular heterogeneity, we plotted Z-scored protein expression values as a heatmap and used epithelial (CK5, CK8, ECAD, Chromogranin A), immune (CD3, CD4, CD11b, CD20, CD45, CD66b, CD68, CD163, FOXP3, HLADRB1, Tryp) and non-immune stromal (αSMA, VIM) markers to create three exclusive categories (Figure S4). This analysis identified 435,702 epithelial cells (62% of cells), 209,521 stromal cells (30% of cells) and 54,238 (8% of cells) immune cells (Figure 2b). We analyzed the percentage of cell subtypes across three clinical grades, which showed an increase in epithelial cells by 21% and 34% and a decrease in immune and non-immune stromal cells by 15% and 27% in high-grade tumors G3 and G4, respectively, compare to TAN samples (Figure 2b).

### The combined expression of epithelial markers shows epigenetic differences between clinical grades

We sought to further define cellular heterogeneity for epithelial, immune and non-immune cells. First, we investigated epithelial cell heterogeneity, by re-clustering only the epithelial cells using the protein markers NCAM, Chromogranin A (ChromA), CD44, VIM, CK5, NKX3.1, AMACR, CK8, ECAD, H3K27ac, H3K4me3, and AR and we identified five epithelial cell clusters: basal, luminal, AMACR-high luminal, neuroendocrine and VIM+ epithelial cells that are likely in the process of epithelial-to-mesenchymal transition (EMT) (Figure 2c).

AMACR is consistently overexpressed in prostate cancer epithelium and is suggested as a cancer-risk associated marker [23]. We observed that 86% of all luminal cells express average levels of AMACR and we found a separate cluster of luminal cells with high AMACR expression in G3 and G4 samples that we labeled as ‘AMACR-high’ (Figure 2c).

While we observed an increase in the percentage of proliferating (Ki67+) AMACR-high luminal cells in G4 samples, the density of Ki67+ AMACR-high luminal cells was similar in G3 and G4 tumors (Figure S4). Significantly, we found an increase in the density of Ki67+ VIM+/EMT cells in G4 tumors, but not in TAN and G3 (Figure S4).

To quantify heterogeneity of epithelial cells, we measured Shannon entropy for each clinical grade [21]. We observed a non-significant decrease in heterogeneity from TAN to G3 and G3 to G4, suggesting epithelial cells in TAN samples show more differences in epithelial marker expression, as opposed to G4 samples (Figure 2d).

Cell subtypes are static categories that may not reflect the heterogeneity of cell states in prostate cancer epithelia. To determine the specific markers associated with the heterogeneity of epithelial cell states, we investigated the combinatorial expression of 12 epithelial markers in 699,461 single-cells (Figure S5). Strikingly, we found that the combined expression of AMACR, H3K27ac, H3K4me3 and AR in epithelial cells showed a grade-specific difference: 21.5% in G4, 6.4% in G3 and 0.4% in TAN (Figure 2e, Figure S5). This grade-specific expression was not observed in CK8+ luminal cells (Figure 2f). We confirmed loss of CK8 in G4 tumors using H&E images of individual TAN, G3 and G4 glands (Figure 2f).

H3K27ac marks active regulatory regions associated with increased transcription [24]. We observed a 10% increase in the number of cells with H3K27ac expression in G4 tumors (Figure S5). To determine whether the spatial distribution of this marker may contribute to disease progression, we calculated Moran’s I values for all epithelial markers (Figure S5) and visualized spatial expression patterns on the tissues (Figure 2g). Moran’s I uses continuous marker expression across a tissue to determine spatial auto-correlation [25]. We observe broad, low-level expression of H3K27ac that overlaps with high-AMACR and AR expression in G4 samples, as opposed to TAN (Figure 2g).

### Identification of mast cell populations that differentially express CD44, CD90 and GZMB

We further stratified immune cells and identified 13 myeloid and lymphoid cell types with Louvain clustering based on the expression of 17 immune markers: CD66b, CD11b, CD20, NCAM, PD1, CD163, HLA-DRB1, CD68, CD90, Tryp, CD44, GZMB, CD45, CD3, CD4, FOXP3 and AR (Figure 3a and Figure S6). Mast cells constituted the second largest population of immune cells (18%) following helper T cells (19%), forming two distinct clusters on the UMAP, which suggested heterogenous mast cell populations in localized prostate tumors (Figure 3a-b).

**Figure 3:**
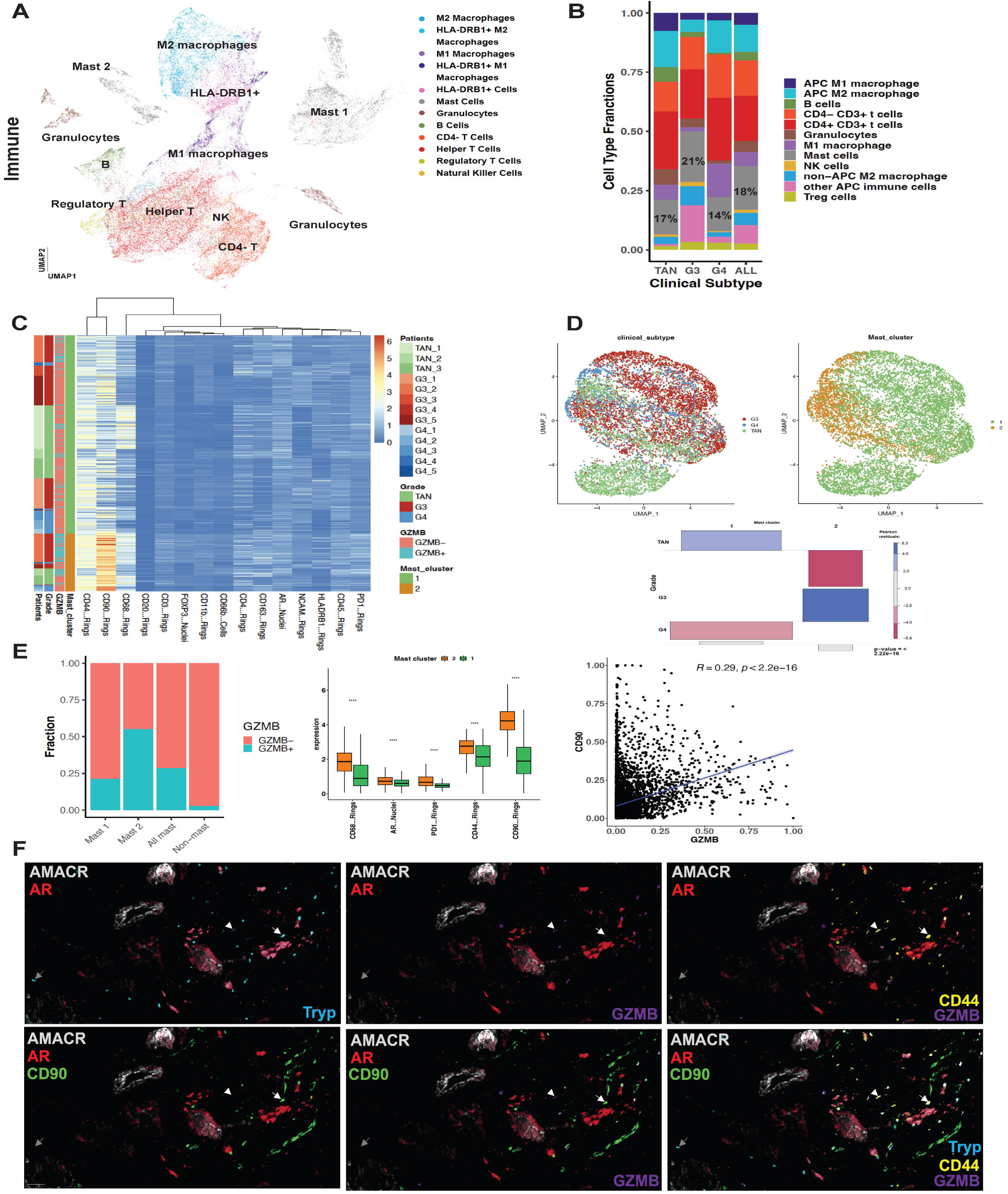
Differential expression of CD44, CD90 and GZMB in mast cells is associated with clinical grade G3. (A) UMAP showing immune cell subtype clusters identified via Louvain clustering. (B) Fractions of immune cell subtypes show high numbers of mast cells across all clinical grades, significantly higher in G3 patients (p<2.22e-16). (C) Heatmap showing log-normalized signal intensity values of each immune marker for the two mast cell clusters identified in our data set, see Mast 1 and Mast 2 clusters in panel A. Large fraction of mast cells shows CD44 expression. (D) Reclustering of mast cells, UMAPs colored by clinical grade (left) and previously identified mast cell clusters (right). Below, in the association plot, the rectangles in each row are positioned relative to a baseline indicating independence. The area of the box is proportional to the difference in observed and expected frequencies. If the observed frequency of a cell is greater than the expected one, the box rises above the baseline, and falls below otherwise. Independence test of mast cell clusters and tumor grades showing significant abundance of mast cluster 2 in G3 patients and mast cluster 1 in TAN patients and no difference in G4 patients. (E) GZMB+/-fraction of mast clusters, mast cells, and non-mast cells (left); boxplot showing protein expression levels differs significantly in mast clusters 1 and 2 (middle); scatter plot showing CD90 and GZMB expression correlation in mast cells (right). (F) An example cycIF image that show heterogenous mast cell populations. White arrow indicates a CD44+CD90+GZMB+AR+ mast cell. Arrowhead indicates a CD44+CD90-GZMB+AR-mast cell. Gray arrow shows a CD44-CD90-mast cell.

Even though the presence of large fractions of mast cells are repeatedly reported in localized prostate tumors, our understanding of their role in mediating tumor response is limited [26]. We determined that the majority of mast cells express CD44, a receptor of hyaluronic acid that is abundant in the extracellular matrix (Figure 3c) [27]. We investigated this further by analyzing differential expression of immune markers in two mast cell populations using Louvain clustering (Figure 3d and Fig 3f).

Mast clusters 1 and 2 showed a striking difference based on the GZMB status. GZMB induces anti-tumor responses and is traditionally found in effector T cells [28]. However, certain GZMB+ cells, such as myeloid derived suppressor cells and mesenchymal stem cells (MSC), can secrete GZMB to degrade receptors on effector T cells and induce immunosuppressive pro-tumor effects [28]. Mast cluster 2 exhibited a higher fraction of GZMB+ mast cells (55%) as opposed to cluster 1 (21%) (Figure 3e). Furthermore, CD90 and GZMB expression in cluster 2 appeared to be correlated with each other (Figure 3c, Figure 3e and Figure 3f). We found a significant increase in the density of proliferating mast cells in high-grade tumors (Figure S6).

### Mast cells with GZMB+ and AR+ expression spatially correlate with M2 macrophages and Tregs in the prostate TME

To determine whether identified mast cell subtypes mediate pro-vs. anti-tumor responses, we examined spatial co-occurrence of mast cells with immune cell types in the prostate TME (Figure 4a). We divided mast cells into two groups based on their GZMB expression and analyzed the frequency of spatial co-occurrence of GZMB+ or GZMB-mast cells with all identified epithelial, immune and non-immune stromal cell types [10]. GZMB+ and GZMB-mast cells did not show any specific enrichment in their spatial auto-correlation in TAN samples (Figure 4b). However, we identified enriched spatial correlation between GZMB+ mast cells and M2 macrophages and between GZMB-mast cells and CD90+ mesenchymal stem cells (MSCs) in G3 tumors (Figure 4c). GZMB+ mast cells also showed highly significant spatial association with M2 macrophages and granulocytes in G4 tumors (Figure 4c and Fig S7). We did not observe GZMB-cell spatial association with MSCs, but with neuronal cells in G4 prostate TME (Figure 4c and Fig S7).

**Figure 4:**
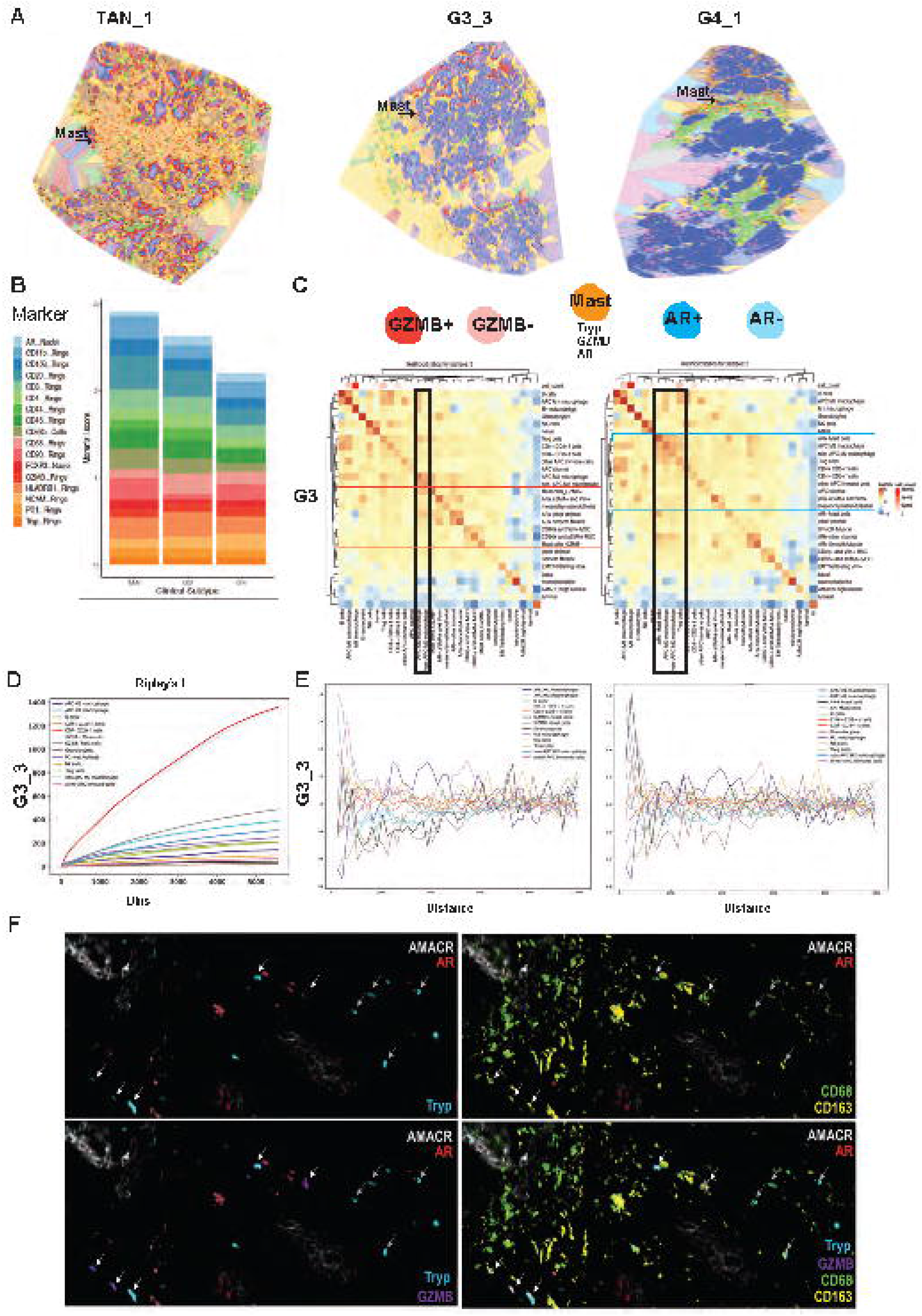
Mast cells spatially interact with distinct immune cell subtypes based on their GZMB status and clinical grade. (A) Voronoi diagrams of neighborhoods for patients TAN_1, G3_3 and G4_1. Blue dots show mast cell distribution across neighborhoods. (B) Average Moran’s I scores of immune cell subtypes across three clinical grades do not show differences based on mast cell marker expression (Tryp) (bottom, orange). (C) Heatmaps of likelihood ratios of direct cell-cell contacts between 27 cell types in G4 tumors where mast cells were divided into two groups as GZMB+/-and AR+/-, respectively. GZMB+ mast cells showed highly enriched spatial correlation with M2 macrophages; and are closely associated with granulocytes. AR+ mast cells showed highly enriched spatial association with Treg cells. (F) Ripley’s L results of patient G3_3 highlight how some cell-types have a more clustered pattern, like CD4+CD3+ T cells (red) and GZMB-mast cells (gray), whereas other have a more dispersed pattern, like Granulocytes (brown) and GZMB+ mast cells (black). (E) Co-occurrence probability of each cell type given the presence of GZMB+ mast cell and AR+ mast cell respectively. The score is computed across increasing distance around each cell in the tissue. GZMB+ mast cells and non-APC M2 macrophage; and AR+ mast cells and CD4-CD3+ T cells show co-occurance in close proximity. (F) cycIF images showing GZMB+ mast cells (white arrows) and M2 macrophage association. Gray arrows show GZMB-mast cells spatially further away from M2 macrophages.

Functional interactions of mast cells with Tregs in colorectal cancer are shown to generate potently immunosuppressive and proinflammatory Tregs [29, 30]. To test whether expression profiles of mast cells impact their spatial correlation, we compared the behavior of mast cells with and without AR expression in the prostate TME. We identified AR+ mast cells that closely interact with TRegs in G3 and G4 prostate tumors as compared to TAN samples (Figure S7). Additionally, AR+ mast cells had enriched spatial association with M2 macrophages in G3, but not in G4 tumors (Figure S7). AR-mast cells did not show any specific enrichment in their spatial association (Figure S7). Taken together, our results show changes in the spatial association of mast cells with specific lymphoid, myeloid and stromal cells based on their GZMB and AR expression, suggesting possible functional roles for distinct mast cell states.

To investigate if there is a direct impact of mast cell clustering versus dispersed mast cells on these spatial correlations, we first generated Voronoi diagrams of each tissue-section and mapped back mast cells on Voronoi neighborhoods (Figure 4a) [10]. Second, we calculated Moran’s I values for each patient (Figure 4b and Figure S7). Our results did not show any differences in mast cell auto-correlation associated with grade (Moran’s I average = 0.08 for all grades), which eliminated the possibility of enriched mast cell correlations mediated by mast cell clustering at the edges of cancer cells. This provides further evidence that the specific spatial association of mast cells with M2 macrophages is not an artifact, but is associated with tumor progression in higher clinical grade tumors (Figure 4b-c).

We validated this result with Ripley’s L statistics similar to Moran’s I using discrete values, cell types [31]. We showed that mast cells did not showed clustering, whereas CD4+ T cells showed high levels of clustering (Figure 4d). We calculated the co-occurrence probability score of each cell type in comparison to GZMB+ and AR+ mast cell populations (Figure 4e). GZMB+ mast cells showed significant spatial co-occurrence with non-APC M2 macrophages and AR+ mast cells showed significant spatial co-occurrence with CD4-cytotoxic T cells based on increasing distance around each cell in the tissue (Figure 4e-f).

### AR+ non-immune stromal cells drive cellular neighborhood organization in tumor samples

We further explored potential predictive factors linked to clinical grade by focusing on non-immune stromal cells. We delineated heterogenous stromal cell populations by clustering them in isolation based on the expression of following markers: αSMA (smooth muscle), VIM (mesenchymal fibroblasts and endothelial cells), CD90 and CD44 (MSCs and cancer-associated fibroblasts), NCAM (nerves), HLA-DRB1 (infiltrating antigen presenting cells) and AR (Figure 5a-b) [32, 33].

**Figure 5:**
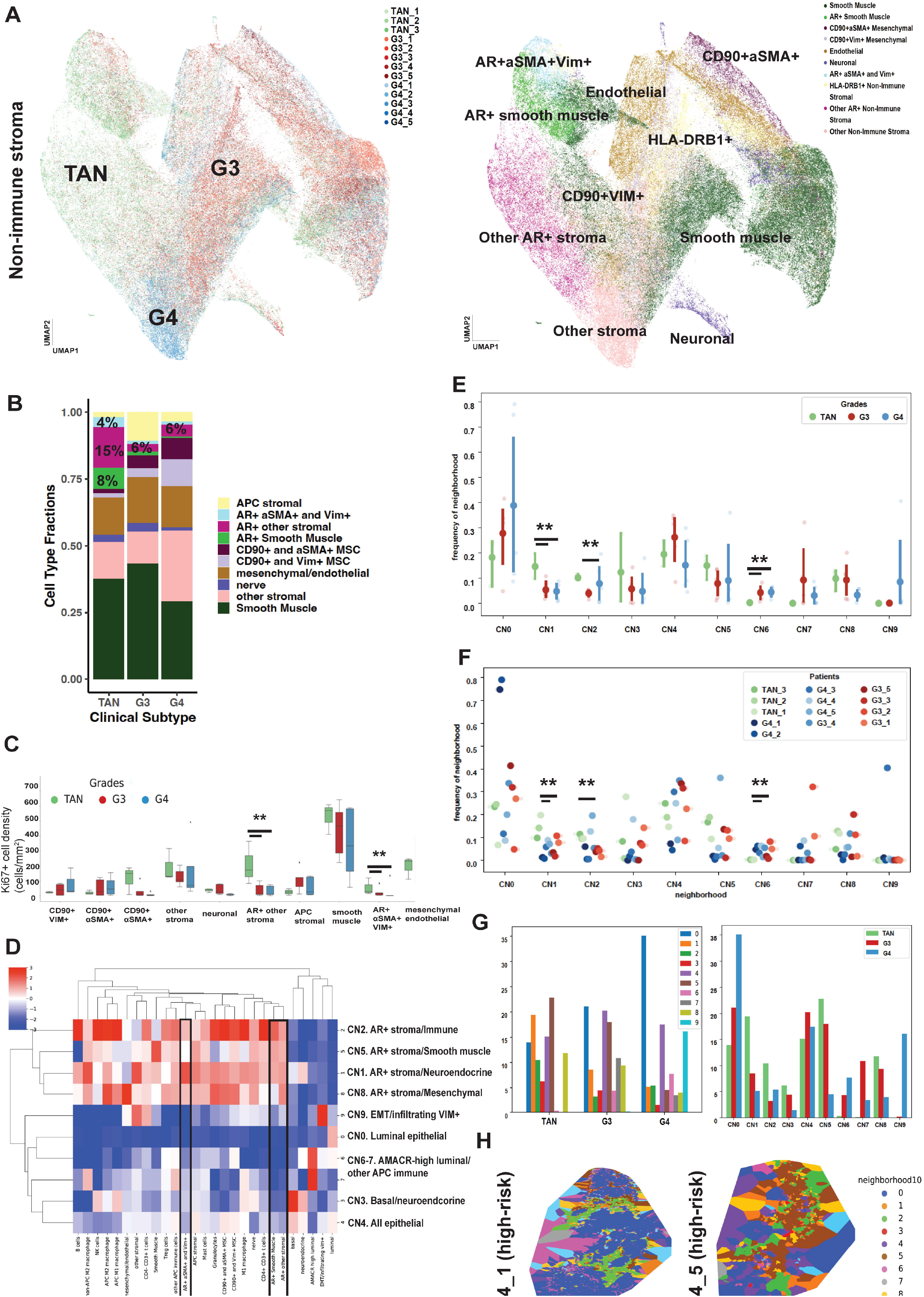
AR positive non-immune stromal cells direct cellular neighborhood organization in TAN vs. G3 and G4 tumor samples. (A) UMAP showing clinical-grade specific (left) non-immune stromal cell subtype (right) clusters identified via Louvain clustering. (B) Fractions of non-immune stromal cell subtypes show high numbers of AR+ stromal cells in TAN samples as compared to tumors. Percentage of CD90+ stromal cells increase in G4 tumors. (C) The density of stromal cell subtypes across three clinical grades shows a significant decrease in the density of two AR+ stromal cell subtypes (AR+ other stromal—p-value<0.03 and AR+ aSMA+/Vim+—p-value<0.04) in prostate tumors compare to TAN. (D) Recurrent cell neighborhoods (CNs) are identified based on 27 cell subtypes. Top four neighborhoods are defined by increased occurrence of AR+ stromal cells, the bottom six neighborhoods are defined by a lack of AR+ stroma. (E) Frequency of each neighborhood is analyzed per clinical grade. CN1 and CN2 (AR+ stromal cells, neuroendocrine and immune subtypes) frequency is significantly higher in TAN as compared to G4 and G3 tumors (t-test p-value 0.017 and 0.016), respectively. CN6 (AMACR-high luminal/other APC immune) frequency is significantly lower in TAN as compared to G4 tumors (t-test p-value 0.001). See Figure S9 for cell type differential enrichment within CNs. (F) Scatter plot showing the frequency of each neighborhood per patient. Each color represents a patient. All G4 patients show enrichment for CNs dominated by epithelial cells except one high-risk G4 patient that is highly enriched for CN5 (M2 macrophages, B cells and AR+ stroma) (G) Bar plots showing the percentages of the Voronoi area of each neighborhood cover in TAN, G3 and G4 samples (right) and actual Voronoi area of each ten CNs per grade (left). (H) Voronoi diagrams of G4_1 and G4_5 patients showing CN distributions. G4_1 and G4_5 patients show enrichment for very different CNs, luminal (blue) vs. AR+ stroma, respectively.

Our results uncover three populations of AR+ stromal cells: AR+ smooth muscle, AR+ αSMA and VIM+ and AR+ other stromal (Figure 5a-b). We determined a significant increase in AR+ stromal cells (27%) in TAN samples, as compared to G3 and G4 tumors (6%) (Figure 5b). Interestingly, despite this increase, we did not observe significant enrichment in the spatial auto-correlation of AR+ stromal cell subtypes in TAN samples, whereas AR+ stromal cells in tumor samples showed significant enrichment in their spatial auto-correlation (Figure S7).

To determine the functional status of these AR+ stromal cells, we examined the density of Ki67+ cell subtypes in the tumor stroma (Figure 5c). The Ki67+ cell density for both AR+ smooth muscle and AR+ other stroma showed a significant decrease in G3 and G4 samples as compared to TAN (p<0.03) (Figure 5c). Combined, these results show the importance of AR regulation in non-immune stroma in the prostate TME.

To determine the impact of AR+ stromal cell populations on tissue organization, we mapped the spatial frequencies of all 27 cell subtypes for each patient and used non-hierarchical clustering to identify recurrent cell neighborhoods (rCNs) [10] (Figure 5d). Each rCN was then mapped back on to the tissue-sections to visualize cell type and neighborhood compositions [10].

Our results indeed show that AR expression in prostate TME strongly informs the 10 rCNs identified across all patient samples. Four neighborhoods were dominated by AR+ non-immune stromal cells interacting with: neuroendocrine (rCN1), mesenchymal/endothelial (rCN8), immune (rCN2) and smooth muscle and M2 macrophages (rCN5). The other six neighborhoods were dominated by the lack of AR+ stroma: luminal epithelial (rCN0), EMT/infiltrating VIM+ (rCN9), AMACR-high luminal cells interacting with APC immune cells (rCN6 and rCN7), basal and neuroendocrine (rCN3) and basal, neuroendocrine and luminal cells (rCN4) (Figure 5d).

We sought to determine if the significant heterogeneity of cell types impact the identified neighborhoods. We calculated the frequency of each rCN for TAN, G3 and G4 samples. Our results identified rCN1 and rCN2, which showed a statistically significant increase in TAN samples (Figure 5e). Both rCN1 and rCN2 are defined by recurrent spatial correlation of AR+ stroma with immune cells (Figure 5d). The frequency of these neighborhoods was significantly lower in both G3 and G4 samples as compared to TAN (Figure 5e-f). We also identified a significant increase in the frequency of rCN6, which consists of AMACR-high luminal cells in G3 and G4 tumors (Figure 5e-f). We repeated neighborhood identification analysis 13 times, leaving one patient out each time (Figure S8 and S9). Our results robustly captured the same rCN composition in each model. AR+ stroma driven TAN vs. G3 and G4 differences were identified in all analyses and AMACR-high driven rCN change was identified in 11 out of 13 (Figure S9).

### Loss of heterogeneity in cellular organization

Our results showed AR+ stromal neighborhoods may distinguish between three clinical grades of prostate samples (Figure 5e). To further delineate these clinical grades, we identified rCNs separately for TAN, G3 and G4 and identified cell types that are differentially enriched within specific neighborhoods across different grades (Figure S8 and S9).

Looking at the frequency of each rCN in three clinical grades, we found TAN samples consist of neighborhoods that have similar frequencies across the tissue-section (less than 25% frequency for each neighborhood) (Figure S8). In contrast, G4 samples consisted of one or two dominant neighborhoods (Figure S8 and Figure 5f). In G3 tumors, the predominant neighborhoods exhibited higher than 25%, but less than 40% frequency (Figure S8). Overall, these results suggest a loss of rCN heterogeneity associated with higher clinical grade, specifically as a result of increase in AMACR-high luminal cells neighborhoods and decrease in AR+ stromal cell neighborhoods (Figure 5f and Figure S8).

We calculated the percentage size distribution of each rCN per clinical grade and showed a decrease in the actual size of the AR+ stromal neighborhoods in G3 and G4 tumors (Figure 5g). The size of the AMACR-high neighborhoods showed an increase in G3 and G4 patients as compared to TANs, supporting our results (Figure 5g).

### Identification of rCNs correlates with patient-specific molecular profiles

To further delineate disease subtypes, we performed low-pass whole genome sequencing for 10 patients, identified common copy number aberrations (CNAs) and analyzed neighborhood differences in G4 tumors. Our results found three dominant rCNs in G4 tumors that had more than 50% frequency. G4_1 and G4_2 tumors exhibited >70% frequency for rCN0 (luminal), which was marked by a significantly smaller number of AR+ smooth muscle cells in G4 tumors as opposed to TAN and G3 (Figure S9). G4_1 patient is a high-risk patient (CAPRA-S=6) and G4_2 is an intermediate-risk (CAPRA-S=3) patient with no known CNAs (Figure S3). G4_3 tumor exhibited >25% frequency for rCN9, which was marked by EMT/infiltrating VIM+ and stromal cells (Figure 5d, Figure S9). rCN9 attracted a significantly smaller number of APC M2 macrophages in G4 samples compared to TAN and G3 (Figure S9). G4_3 patient is an intermediate-risk patient (CAPRA-S=5) with four known amplification events: NCOA2, NBN and EGFR. G4_4 tumor exhibited >30% frequency for rCN3 and rCN4, which were marked by luminal and neuroendocrine epithelial cells (Figure 5d, Figure S9). G4_4 patient is an intermediate-risk patient (CAPRA-S=4) with three known deletion events: PTEN, CDH1 and NKX3.1 (Figure S3). PTEN loss is associated with an immunosuppressive microenvironment that is mediated by TRegs and M2 macrophages [34], which correlates with our results. Finally, G4_5 tumor exhibited >50% frequency for rCN5, which was marked by M2 macrophages and AR+ stroma (Figure 5d, Figure 5g, Figure S9). G4_5 patient is a high-risk patient (CAPRA-S=9) with multiple amplification (PIK3CA and EGFR) and deletion (PTEN, CHD1, RB1, FOXO1, TP53, CDKN1B and NKX3.1) events.

Overall, our combined genomic profiling and cycIF imaging results further stratify G4 tumor subtypes and exhibit major differences in TME organization that may be associated with known CNAs in prostate cancer, underlining the need for understanding patient-specific trajectories for precise treatment approaches (Figure 5h).

### AR+ stromal cells in localized prostate tumors show activated NF κB signaling

To gain further understanding of the biological targets and pathways in AR+ stromal cells, we shifted from imaging and focused on identifying potential gene networks active in AR+ cells in prostate TME. We performed sci-ATAC-seq on flash-frozen tumors collected from the same patients [17] and identified myeloid, lymphoid and non-immune stromal cells based on cisTopic clustering and gene expression scores [35, 36] (Figure 6a). We identified differentially accessible regions (DARs) of AR+ vs. AR-null (AR0) cells and observed significantly enriched accessibility in chromatin regions that regulate NF κB signaling in AR+ stromal cells (Figure 6b) [37]. GO analysis was performed using GREAT and the Panther classification system [37]. To gain more mechanistic insights, we investigated transcription factor (TF) binding motifs significantly enriched in AR+ vs. AR0 stromal cells using both Cistrome and JASPAR [38] (Figure 6c). We identified significantly enriched TFs that are known AR regulators in AR+ cells such as SP1, YY1 and NFYA (Figure 6d). Interestingly, we observed AR+ stromal cells in localized prostate tumors showed significant enrichment for the hypoxia marker HIF1α, suggesting a role for stromal AR regulation in hypoxia. We did not observe any enrichment for CXCR4, a known inflammatory marker, nor for FOXA1, a known repressor of HIF1α (Figure 6e). While there was no significant difference in IFNγ accessibility, a subgroup of AR0 cells were accessible for IFNγ, as expected (Figure 6d). Finally, we confirmed that NF κB1 was significantly more accessible in AR+ cells in the prostate TME (Figure 6d), validating our GREAT GO results (Figure 6b). Overall, our sci-ATAC-seq analysis identified active transcriptional networks in both AR+ and AR0 stromal cells and defined utilizing markers that can be used to further investigate AR+ stromal cells in cycIF imaging.

**Figure 6:**
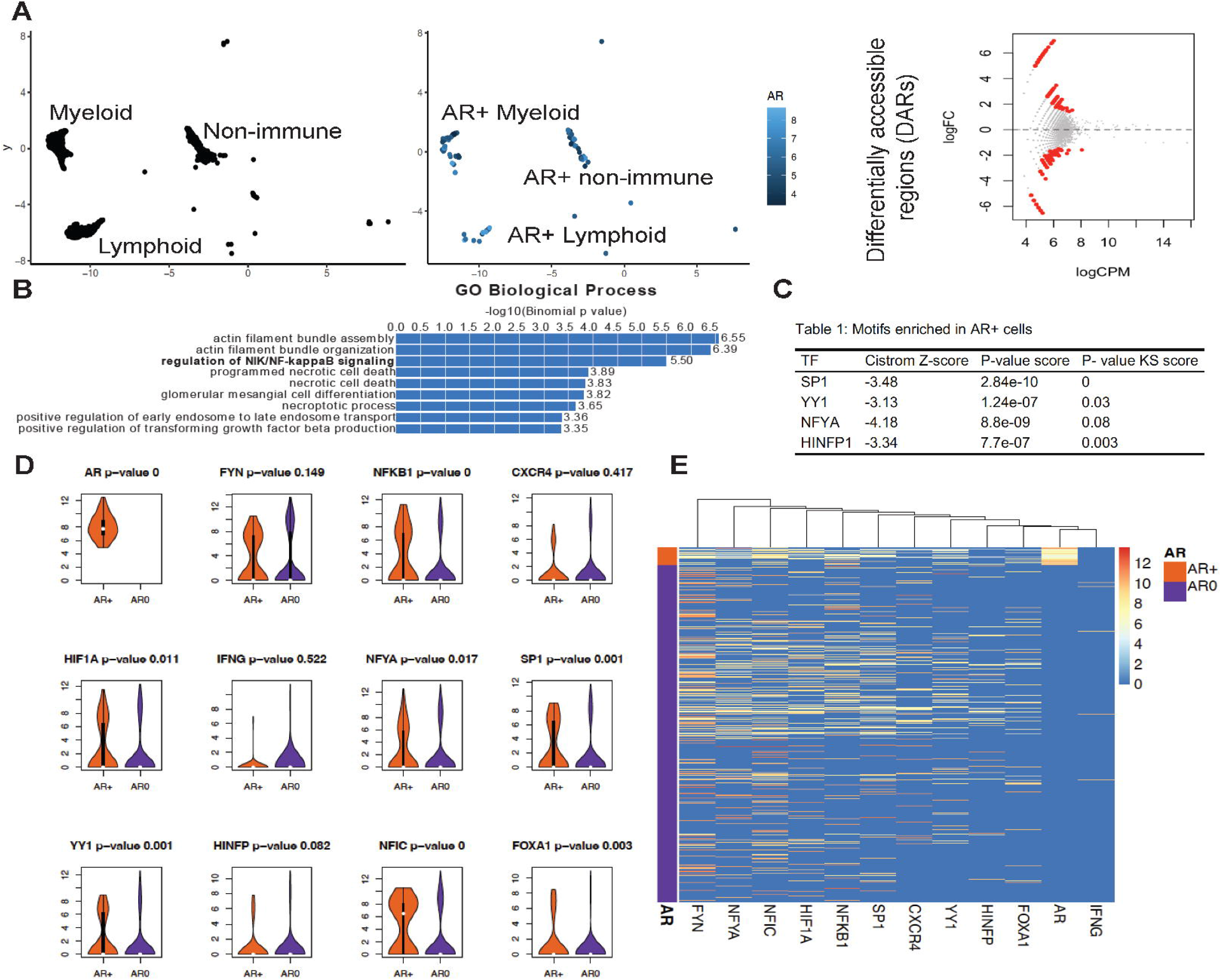
Transcriptional network of AR+ stromal cells in localized prostate tumors show an activation of NF κB signaling and hypoxia. (A) AR+ myeloid, lymphoid and non-immune stromal cells were identified in single-cell ATAC sequencing data from the same patients. Differentially accessible regions (DARs) are identified in AR+ vs. AR0 cells. (B) GREAT GO results show an enrichment for NIK/NF κB signaling pathway regulators in AR+ stromal cells. (C) TF motif binding sites identified in AR+ stromal cells show an enrichment for known AR regulators and HIF1α. (D) Gene scores of identified TFs and pathways in single-cell ATAC sequencing data. (E) Heatmap showing fold enrichment values for differentially accessible regions in AR+ vs. AR0 stromal cells identify significant HIF1α and NF κB accessibility in AR+ cells.

## DISCUSSION

In this work, we integrated multiple single-cell approaches with bulk genomic sequencing and pathology to investigate changes in the prostate TME using patient samples across three clinical grades. We present novel spatial patterns within localized prostate tumors that imply epigenetic and molecular disease subtypes even within tumors of seemingly identical pathological grade. While 13 specimens are not representative of all known molecular subtypes of localized prostate cancer, nor do they allow for full statistical analyses, our results show striking differences in disease subtypes within a clinical grade, which would not be detected based on an H&E image. Future studies will involve further delineation of disease subtypes by generating an increased number of multi-modal spatial maps and examining their predictive and prognostic clinical utility.

Our multi-modal analyses underline the need for generating more detailed disease subtypes for patients with localized prostate cancer to improve prognosis. Biological underpinnings measured for each G4 tumor exhibit differences in CNAs, immune response and neighborhood organization. For instance, while we observed an overall increase in the frequency of immune-cold luminal neighborhoods in G4 tumors, we identified a G4 patient dominated by an M2 macrophage-AR+ stromal cell neighborhood. Interestingly, this patient was the only patient in our cohort with an RB1 deletion, which is a common event in prostatic small cell carcinoma [39].

We also identified similarities between patients for each clinical grade that may be used for automated patient stratification in future studies. We show the combined expression of AMACR, H3K27ac, H3K4me3 and AR in epithelial cells can distinguish between TAN, G3 and G4, independent of CK8 status. Furthermore, we report a loss of heterogeneity in high-grade prostate tumors as compared to low-grade and TAN samples. Our first line of evidence for loss of heterogeneity comes from our previous single-cell epigenomic analyses that demonstrate increased similarity in chromatin accessibility profiles of high-grade tumors [17]. Here, we observed increased epithelial expression of epigenetic markers H3K27ac and H3K4me3, which are associated with activated transcription and accessible chromatin, accompanied by a decrease in Shannon entropy in high-grade tumors. Additionally, using cycIF and spatial statistics, we observed the specific areas of increased similarity in cancer cells on patient-sections, identifying overlapping expression of H3K27ac and AMACR in G4 tumors. Finally, we report patient-specific rCNs in high-grade prostate tumors, showing that decreased heterogeneity is also reflected at the tissue-organization level.

As shown in previous work [18, 40], the use of whole-tissue sections instead of TMAs allowed us to capture long-distance associations across 27 cell subtypes. We observed changes in spatial association of mast cell populations with M2 macrophages and TRegs based on their GZMB and AR status. Mast cells are a suggested independent prognostic variable for prostate cancer progression [26]. CD44 expression is shown to be a critical factor for mast cell proliferation, differentiation and maturation in other diseases [27]. Here, we report large fractions of mast cells expressing CD44, with no grade-specific changes in localized prostate tumors. It is possible that CD44 expression in mast cells modulates mast cell attachment to the extracellular matrix and contributes to ECM reorganization in high-grade tumors. Detailed markers are needed to identify CD44 isoforms that may be heterogeneously active across clinical grades. Furthermore, we identify specific spatial associations between mast cells that express GZMB and AR and M2 macrophages and Tregs. Functional interactions between mast cells and Tregs and macrophages mediated by cytokines have been investigated in mouse models of colorectal cancer [29, 30]. The interaction between mast cells and Tregs lead to potently immune suppressive and proinflammatory Tregs. Mouse models of prostate cancer cells do not have abundant mast cell populations but we show these mast cell populations may play a role in an immunosuppressive behavior in human adenocarcinoma. Co-culture studies are needed to investigate the functional interactions between GZMB+ and AR+ mast cell populations with M2 macrophages.

Loss of stromal AR in advanced prostate cancer is associated with biochemical relapse and poor prognosis [16]. However, there are no previous reports of delineating non-immune stromal cell types in human adenocarcinoma based on AR expression. While we detected 10 distinct stromal cell populations in this study, we observed many unclassified non-immune stromal cells that were marked positive for AR. We report that this ‘AR+ other stroma’ cell population played a major role in neighborhood organization across three clinical-grades. We suspect there will be tremendous value in stratifying these ‘AR+ other stroma’ cells to better understand their role in neighborhood organization in low-vs. high-risk patients and in response to AR inhibitors. In future studies, additional markers will be added to cycIF panels to specifically stratify AR+ stromal cells, such as markers identified through sci-ATAC-seq.

sci-ATAC-seq provided valuable insights into active pathways in AR+ stromal cells. The number of cells captured via sequencing remains low as compared to the number of single-cells captured via imaging. However, the depth of cis-and trans-regulator information gained from sci-ATAC-seq is unmatched in multiplex imaging, which is limited to the number of protein markers that can be probed from a single tissue-section. Integrating single-cell sequencing and imaging approaches allowed us to overcome limitations of each experimental tool and unlocked mechanistic insights related to NF κB and hypoxia activation in AR+ stromal cells. We propose the use of multi-modal single-cell approaches to understand the role of stromal AR expression in regulating inflammation.

## METHODS

### Clinical samples

Thirteen patients with treatment-naïve, localized prostate cancer were identified from the OHSU’s Biolibrary using medical records and reviews of H&E images. Five unstained 5-micron FFPE adjacent tissue sections and an adjacent H&E stained tissue section from each patient were obtained. H&E images were reviewed by a pathologist and a Gleason score was assigned to each patient. Patients were classified into risk groups based on the The Cancer of the Prostate Risk Assessment Post-Surgical (CAPRA-S) scoring system (Figure S3) [41].

### Cyclic IF Antibody Panel

Antibodies used in the study are shown in Supplementary File 1. Tissue sections used in antibody validations are shown in Supplementary File 2. Antibodies were either obtained as commercially conjugated to fluorophores or in a BSA and Azide free buffer and conjugated to AlexaFluor (AF) 488, AF555, AF647 and AF750 dyes in-house (ThermoFisher Scientific, A20006) from as previously described [21].

### Fluorophore conjugation to antibodies

Antibodies were either obtained as commercially conjugated to fluorophores or in a BSA and Azide free buffer and conjugated to AlexaFluor (AF) 488, AF555, AF647 and AF750 dyes in-house. AlexaFluor NHS ester reagents (ThermoFisher Scientific, A20006 and A20000) were dissolved in DMSO to a final concentration of 10 mM. Antibodies were reacted with fluorophore dyes at a 1:10 antibody to dye molar ratio in 0.1 M sodium bicarbonate buffer of pH 8.3. Conjugation reactions were carried out on a rocker for two hours at room temperature in the dark. A buffer exchange step using 10 kDa Amicon Ultra spin columns (Sigma-Aldrich, UFC5010) was performed to remove the excess Alexa Fluor dye from conjugation reactions.

### FFPE tissue processing

FFPE tissue sections were deparaffinized as previously described [17]. The paraffin embedding was removed from the tissue sections via an overnight incubation at 50^°^C followed by a one-hour incubation at 65^°^C. Slides were immediately transferred into Xylene and sequentially immersed in a fresh Xylene solution two times, 5 minutes each; 100% ethanol two times 5 minutes each; 95% ethanol two times 2 minutes each; 70% ethanol two times 2 minutes each and distilled water two times 5 minutes each. Antigen retrieval was done in a medical decloaking chamber (BioSB, TintoRetriever Pressure Cooker).Tissue slides were placed in a Coplin jar containing 10mM citrate buffer, pH 6 (Sigma-Aldrich, C-1909) and incubated at high pressure for 15 minutes. Each slide was then immersed in hot distilled water and transferred into 1X Target Retrieval Solution, pH 9 (Agilent, S2367) for 15 minutes. Following antigen retrieval, tissues were briefly washed in ddH2O, followed by 5 minutes in 1X phosphate-buffered saline (PBS), pH 7.4 (Fisher, BP39920).

### Cyclic IF imaging of prostate tissues

Cyclic IF protocol was adapted as previously described with minor revisions [42, 43]. Prior to immunofluorescence staining, the slides were incubated in a quenching solution (10% 10X PBS, 0.4% 5M NaOH, 3% H2O2) under broad spectrum light for 1 hour to reduce inherent tissue autofluorescence. After quenching, tissues were blocked with 5% NGS (Fisher Scientific, S-1000) in 1% BSA (Thermo Scientific, AM2616) solution for 30 minutes at room temperature. For each round of cyclic IF staining, primary antibodies were diluted in 5% NGS in 1% BSA. Slides were stained overnight at 4^°^C in a humid chamber. Antibodies were washed four times in 1X PBS for 5 minutes. Tissue sections were cover slipped using Slowfade Gold DAPI mounting media using a #1.5 thickness coverslip (Corning Life Sciences, 2980-243 and 2980-245) and imaged using a Zeiss Axio Scan.Z1 slide scanner. After a successful scan was obtained, the fluorophore signal was quenched as described above under broad spectrum light for 1 hour. Slides were washed 3 times in 1X PBS for 5 minutes. Quenching was confirmed under a microscope. Primary antibody staining, imaging and quenching steps were repeated for 10 rounds, using up to four distinct antibodies conjugated to AF488, AF555, AF647 and AF750 and DAPI nuclear stain in each round. The order at which each antibody was used in the cyclic IF experiment can be found in Supplementary Figure 2B. Quenched signal images were taken following rounds 3 and 10 and prior to round 1.

### Whole slide scanning

An Axio Scan.Z1 microscope (Zeiss, Germany) with a Colibri7 LED light source with Cy7 line (Zeiss, Germany) was used for automated high-throughput whole slide scanning. LED intensities were kept constant for all rounds of imaging, where the LED exposure times was adjusted in each round. Exposure times were kept constant between all tissues in each round. Filter sets for DAPI, GFP, Cy3, Cy5 and Cy7 were used for 5 channel imaging in each round of cyclic IF. Images were taken with an Orca Flash 4.0 V3 CMOS camera (Hamamatsu Photonics, Japan) at 20X resolution using a plan-apochromat 20x/0.8 objective (Zeiss, Germany). Image acquisition profiles were set using the ZEN Blue edition imaging software, where the coarse and fine focus strategies and light exposure times were manually adjusted for each round of cyclic IF imaging.

### Low-pass WGS library preparation, sequencing and data analysis

Low-pass WGS libraries were created using the Kapa Biosystems Hyper Prep kit (KAPA Biosystems, Capetown, South Africa) using 11.8 ng/ml of G4_1; 17.6 ng/ml of G4_3; 13.8 ng/ml of G4_4; G4_4 10.8 ng/ml; 13 ng/ml G4_5; 13.3 ng/ml G3_1; 9.2 ng/ml G3_2; 11.4 ng/ml G3_3; 9.6; G3_5; 11 ng/ml TAN_1; 10.1 ng/ml TAN_2; 10.4 ng/ml TAN_3 DNA as input. DNA was fragmented by sonication to 150 bp using a Covaris E220 (Covaris Inc., Woburn, MA, USA) and ligated to dual-indexed Illumina-compatible sequencing adaptors using KAPA Hyper Prep Reagents and protocol and then PCR amplified for 9 cycles using Illumina library amplification primers. Amplified libraries were assessed for size distribution and concentration using the 2100 Bioanalyser system (Agilent Technologies, Santa Clara, CA, USA) with HS dsDNA Kits. Samples were then sequenced the Illumina NextSeq 500, paired-end 75 bp with dual 14-bp indexing cycles. Raw fastq files were demultiplexed using in-house software, aligned using BWA-MEM, and copy number alterations were called using the IchorCNA software package using default parameters [44] (https://github.com/broadinstitute/ichorCNA).

### Single-cell ATAC sequencing

#### Sample Preparation

Radical prostatectomy samples from 13 patients with low-and high-risk prostate cancer were obtained as flash frozen samples. Samples were prepared for single-cell ATAC sequencing using a combinatorial indexing platform as previously described [17, 45]. Tissues were dounce homogenized five times using the loose and tight pestles in Nuclei Isolation Buffer (NIB, 10□mM TrisHCl pH7.4, 10□mM NaCl, 3□mM MgCl2, 0.1% IGEPAL, 1 protease inhibitor tablet (Roche, Cat. 11873580001)). After nuclei isolation, the sample was centrifuged at 500 x g for 5 min at 4 °C and the pellet was washed with 1XPBS with another 2X centrifugation steps. Nuclei were resuspended in NIB and a single nuclei suspension was prepared by filtration through a 35 mm strainer (Corning).

#### Combinatorial Indexing for single-cell ATAC sequencing

Nuclei were stained using DAPI and 3000 nuclei were sorted in each well of a 96 well plate using a flow sorter (BD FacsAria Fusion cell sorter FACSDiva v8.0.3). Each well contained 5□μL NIB and 5□μL TD buffer from Illumina, and 1 mL of 2.5 mM uniquely indexed transposome. Tagmentation was carried out at 55 °C for 30 minutes, plate was placed on ice following tagmentation. Nuclei from each well were pooled and passed through a 35 mm filter and re-stained with DAPI. Pooled nuclei were loaded into the FacsAria flow sorter and 22-25 nuclei were sorted into each well of another 96-well plate. Wells contained a master mix to facilitate denaturation and PCR steps that followed (0.25□μL 20□mg/ml BSA, 0.5□μL 1% SDS, 7.5□μL distilled water and 2.5□μL i5 and i7 PCR index primers (IDT)). Plate was incubated at 55 °C for 15 minutes to denature the transposase. PCR master mix containing NPM PCR mix (Illumina), 1X SYBR Green dye and uniquely indexed PCR primers were added to each well. Fragmented DNA was amplified in RT-PCR (QuantStudio v1.7.1) with the following set-up: 75□°C for 5□minutes, 98□°C for 30□seconds, (for 22–25 cycles) 98□°C for 10□seconds, 63□°C for 30□seconds, 72□°C for 60□seconds, plate read at 72□°C for 10□seconds. qPCR cycles were monitored and amplification was stopped prior to saturation.

#### sci-ATAC-seq library preparation

DNA from all wells were pooled and purified using Qiagen PCR purification kit followed by size selection with AMPure beads. DNA was quantified and fragment size distributions were visualized on a Bioanalyzer. Libraries were sequenced using NextSeq 500 platform (Illumina, NextSeq500 NCS v4.0) using a 150-cycle kit with a custom sequencing recipe (Read 1: 47 cycles; Index 1: 8 imaged cycles, 27 dark cycles, 10 imaged cycles; Index 2: 8 imaged cycles, 21 dark cycles, 10 imaged cycles; Read 2: 47 cycles).

#### Data analysis with snapATAC

Sequencing base calls were converted to fastq files using bcl2fastq. Reads were pre-processd using snaptools *snap-pre* command with default parameters and aligned to the hg19 genome using snaptools *align paired-end* command using default parameters and *bwa* specified as the *aligner* [36]. Peaks were called using *runMACS* command from snapATAC R package with the following MACS2 parameters: *-nomodel -shift 100 -ext 200 -qval 5e-2 -B –SPMR* [36]. Peaks called from each sample individually, each cluster individually and the whole dataset were merged into a master peak matrix consisting of 125,569 peaks used for downstream analysis.

The number of transposase insertion events were quantified in 5 kb bins in the genome to generate a binned counts matrix using the snaptools *snap-add-bmat* command. This matrix was binarized and input into a latent dirichlet allocation dimensionality reduction using the snapATAC function “*runLDA”* using 30 topics. Clusters were identified using the “*runCluster”* function which utilizes the Louvain method for community network detection. Clusters consisting of immune or non-immune stromal cells were identified based on cluster IDs and gene expression scores [17, 36].

#### Identify Differentially Accessible Regions (DARs)

We used the “*findDAR”* function from *snapATAC* R package *edgeR* (v3.18.1) with BCV=0.4 for humans, which takes a snap object and finds differentially accessible regions (DARs). P-value was then adjusted into False Discovery Rate (FDR) using Benjamini-Hochberg correction. We identified AR+ and AR null single-cells based on chromatin accessibility near AR gene [36]. Peaks with FDR less than 0.05 were selected as significant DARs. We ranked the elements based on their enrichment p-value and picked up the top 2,000 peaks for motif analysis.

#### Motif analysis

We used the function “*getJasparMotifs”* from R package *chromVar* that fetches motifs from the JASPAR database [46]. The function “*matchMotifs”* from the *motifmatchr* R package finds which peaks contain which motifs (Schep, 2022). We found enriched motifs of both AR+ and AR0 cell groups using their DARs. One option is the p.cutoff for determing the stringency of motif calling. The default value is set to 0.00005, that gives reasonable numbers of motif matches for human motifs. The function can also return the number of motif matches per peak and the maximum motif score per peak. We calculated enrichment p-values comparing motif scores per peak between AR+ and AR0 cell groups using Kolmogorov-Smirnov test implementation of *stats* library in R (R Core Team, 2013).

### Cyclic IF image analysis pipeline

#### Image Preprocessing

All images were acquired as .czi files from the Axio Scan Z1 microscope (Zeiss, Germany). ZEN Blue image processing software (Zeiss, Germany) was used to convert .czi images to 64Bit BigTiff images without any compression of the original grayscale data for each channel. The following steps; registration, segmentation and autofluorescence subtraction, were performed.

A custom image registration algorithm was used to overlay DAPI images from each round of cyclic IF imaging^107^. Registration quality was confirmed by visualizing on QiTissue software (Quantitative Imaging Systems, LLC, Pittsburgh, Pennsylvania) or Napari application in Python^109^.

#### Background Signal Removal

Images acquired after the chemical quenching step following the third round of cyclIF experiment were selected as the autofluorescence image. The autofluorescence image was subtracted from each round. Following autofluorescence subtractions, a background smoothing operation was performed to smooth out staining gradients and uneven illumination patterns on the images. This was done by applying white top hat operation with the structuring element radius equal to the biggest nucleus radius within each tissue.

#### Image Segmentation

Cellpose was used for the segmentation of individual nuclei [20]. For nuclei segmentation, the z-projection image of all DAPI images from each round of cyclic IF imaging was used. Cytoplasm segmentation mask was created by dilating the nuclear segmentation mask. Average signal intensity values from individual nucleus, cell membrane, cytoplasm (nucleus minus cell membrane) and whole cell (nucleus plus cell membrane) compartments were calculated.

To eliminate parts of tissue that degraded and fell off the slides, we created a binary mask from the DAPI image from the last round of imaging and determined and used only the unique cell labels present in the last round DAPI mask.

#### Signal Intensity Normalization and Marker Thresholding

Signal intensities of each marker were normalized by background fluorescence intensity. A multi-step approach was followed for background fluorescence determination. First, we used an automated algorithm called RESTORE, which uses mutually exclusive markers, to calculate background fluorescence intensities [47]. Cutoff value outputs from RESTORE were visually inspected for each marker in each tissue and manually adjusted. Cutoff values were then used to normalize signal intensities by dividing the raw fluorescence intensity values by the cutoff value. Therefore, values above “1” were “marker positive” as their signal intensity values were above the background signal intensity. Based on normalized intensity values, each cell in the dataset was labeled as proliferating (Ki67+) or non-proliferating (Ki67-) using a threshold of normalized Ki67 intensity >1. Similarly, each immune cell was identified as cytotoxic (GZMB+) or non-cytotoxic (GZMB-) and AR expressing (AR+) or not (AR-).

#### Clustering and cell type identification

Cell type identification analysis was performed as previously described using publicly available Python and R packages [21]. Briefly, single-cell phenotypes were identified based on the normalized signal intensity values from our 28-plex cyclic IF data. To cluster single-cells, we used an unsupervised Louvain community detection algorithm called PhenoGraph (k=40, to detect rare cell phenotypes) [48]. Prior to the application of PhenoGraph to our data, we performed 99^th^ percentile normalization and arcsin transformation (cofactor=5). The GPU-accelerated implementation of PhenoGraph was implemented from https://gitlab.com/eburling/grapheno [21]. For heatmap visualization of single-cell phenotypes and respective marker expression, “*clustermap”* function from seaborn package in Python was used from previously developed algorithms available at https://gitlab.com/eburling/BCTMA/-/blob/master/pyviz_figs.py.

Following PhenoGraph, clusters were visually confirmed by mapping the location of cells from each cluster onto segmentation masks. We selected random samples to validate cluster annotations by doing side-by-side comparisons of cells mapped onto segmentation masks and IF images. MATLAB code for mapping cells onto segmentation masks is available on GitHub (https://github.com/zeynepsayar/PCa_immune/blob/main/mapping.m).

#### UMAP Visualization

To enable the 2-dimensional visualization of our high-plex feature-by-cell data, we used the GPU accelerated, RAPIDS library adaptation of UMAP on 99^th^ percentile normalized and arcsin transformed (cofactor=5) data. The default set of parameters were used for both visualization methods.

#### Quantification of epithelial marker heterogeneity using Shannon Entropy

Cells from each tissue were labeled as positive for CK8, CK5, AR, AMACR, H3K27ac, H3K4me3 and NKX3.1 if their normalized signal intensities were above 1. A Shannon entropy figure (Figure 2D, Supplementary Figure 5) was generated using UpSetPlot as implemented in the following algorithm: https://gitlab.com/eburling/BCTMA/-/blob/master/rapids015_figs.py. The heterogeneity of epithelial markers was computed by measuring the Shannon entropy [21]. Statistical differences between marker heterogeneity in TAN, G3 and G4 clinical grades were quantified using the pairwise_tukey function from pingouin [49].

#### Combined marker expression analysis

To test the significance of the association between clinical grade and combined expression of epithelial markers (AR, AMACR, H3K27ac, H3K4me3) we used mosaic plots. Mosaic plots visualize the deviance of observed combined marker expression in each clinical grade from expected combined expression frequencies of an independence model. Mosaic plot in Figure 2E was generated with “*mosaic*” function from R’s “*vcd*” package using default parameters. The size of boxes in mosaic plot are proportional to the difference between expected and observed frequencies. Boxes are colored in red if observed frequencies smaller than expected, and blue if observed frequencies are greater than expected. We performed a chi-square test with the null hypothesis that grades and combined marker expressions are independent. P-values < 0.001 reject the null hypothesis and indicate grades and combined marker expressions are dependent.

#### Mast cell differential marker expression analysis

Normalized mean intensity values of CD68, AR, PD1, CD44 and CD90 were plotted for each cell in Mast cluster 1 and Mast cluster 2 from all samples in the cohort. t-test was used to identify any statistically significant differences between individual and combined marker expressions between Mast cluster 1 vs. 2. p-values < 0.05 were significant.

#### Cell density calculations

Cell counts were calculated for each cluster in epithelial, stromal and immune populations. Tissue areas were calculated in mm^2^ using ImageJ software (NIH, Bethesda, Maryland). Whole tissue region was selected as the region or interest (ROI) and the area was calculated in mm^2^ using the measure option and the manually set scale. The fraction of Ki67-positive proliferating cells within each immune, epithelial and stromal cell subtypes was calculated and plotted in Figure 5E. Frequencies of each cell phenotype found in each tissue section were plotted in Figure 3B. Statistical significance of differences between cell densities were tested using a t-test (p<0.05).

#### Spatial analysis

#### Pairwise cell-cell interactions

Pairwise cell-cell interactions were calculated using the R Shiny app “app_CRC_contacts.R” as previously described (https://github.com/nolanlab/NeighborhoodCoordination) [50]. Cell types were omitted if the number of unique adjacent cells for that cell type was less than 100 cells. Calculated values were mapped using the *ComplexHeatmap* package in R. To calculate pairwise cell-cell contacts, we calculated likelihood ratios of the 27 cell types that we identified as follows:

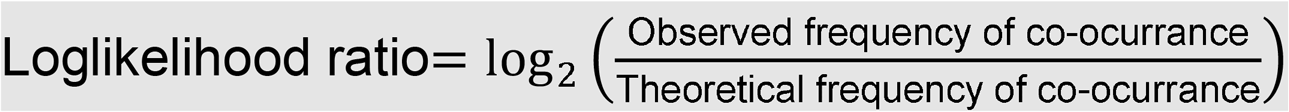

Voronoi diagrams were generated through the following process in https://github.com/nolanlab/NeighborhoodCoordination: FCS files obtained from VorteX were exported and then processed using a custom Java algorithm. This algorithm was designed specifically to create Voronoi diagrams and derive contact matrices between individual cells.

#### Cell Neighborhood identification

Cell neighborhoods were determined by using existing analysis platforms found in https://github.com/nolanlab/NeighborhoodCoordination. The ten nearest neighbors of each individual cell were determined. Each window was represented as the frequency of each cell type found within the ten nearest neighbors. In total we identified 27 cell types, therefore each window was a vector of length 27. To determine the cell neighborhoods, these windows were clustered using K-means clustering with k=5 and k=10 with Python’s *scikit-learn* library and “*MiniBatchK-Means”* function. We identified significantly enriched neighborhoods in each clinical grade by performing t-tests (p<0.05).

To validate the reproducibility of cell neighborhoods identified from the dataset and their enrichment in specific clinical grades, we iteratively performed cell neighborhood analysis by leaving out one patient at a time. For example, we identified cell neighborhoods from all cells in the dataset excluding cells from patient 1 and performed t-test to identify which cell neighborhoods were significantly enriched in TAN vs. G3 vs. G4. We repeated this analysis for each patient in the cohort.

#### Enrichment of cell types within cell neighborhoods

To calculate differential cell type enrichment in rCNs, differential enrichment analyses using linear models was conducted. The models were estimated using the following equation: Yn,c = β0 + β1X + β3Yc + e. In this equation, Yc represents the logarithm of the overall frequency of cell type c, X is an indicator variable for the patient group, Yn,c represents the logarithm of the frequency of cell type c in CN n, βi are coefficients, and e represents Gaussian noise with a mean of zero. To avoid computational issues, a pseudocount of 1e-3 was added before taking logarithms. The estimation of these models was performed using the statsmodels Python package. The resulting coefficient estimates and p-values for β1 were extracted and visually presented (significant if p<0.05).

#### Spatially variable markers with spatial autocorrelation statistics (Moran’s I Score)

Moran’s I evaluates whether a marker is clustered, dispersed or random based on the spatial expression of that marker:

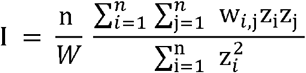

where *z_i_* is the deviation of the feature from the mean 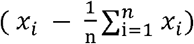, *w_i,j_* is the spatial weight between observations, *n* is the number of spatial units and *W* is the sum of all *w_i,j_*.

The Moran’s I global spatial auto-correlation statistics were calculated using Python’s *Squidpy* package implementation [31]. We computed the Moran’s I score with “*squidpy*.*gr*.*spatial_autocorr*” and *mode = ‘moran’*. We first generated a spatial graph with “*squidpy*.*gr*.*spatial_neighbors*” which provides a score on the degree of spatial variability of marker expression. The statistic as well as the p-value were computed for each marker, and FDR correction was performed (significant if p< 0.05). We visualized the top markers after applying the Moran’s I spatial autocorrelation statistics.

#### Spatially variable cell-types with Ripley’s L (Ripley’s spatial statistics)

Ripley’s spatial statistics is a family of spatial analysis methods used to describe whether points with discrete annotation (i.e., cell-types) in an area of interest follow random, dispersed or clustered patterns. To calculate Ripley’s L, we used “*squidpy*.*gr*.*ripley”* function with *mode = ‘L’*.

#### Cell-type co-occurrence ratio

Co-occurrence ratio of cell-types provides a score on the co-occurrence of the cell-type of interest across spatial dimensions. It is defined as follows:

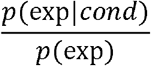

Where p(exp|cond) is the conditional probability of observing a cell-type exp conditioned on the presence of a cell-type cond, whereas p(exp) is the probability of observing exp in the radius size of interest. The score was computed across increasing radii size around each cell in the tissue. We used Python’s “squidpy” library, “*squidpy*.*gr*.*co_occurrence”* function to calculate cell-type co-occurrence ratio [31].

## Author Contributions

ZS conducted cyclic IF experiments. CA and ZS analyzed the cyclic IF imaging data. CA and AC analyzed single-cell ATAC-seq data. CB and SEE performed CNA experiments and analysis. GT reviewed pathology slides. RK and VS reviewed clinical patient data. ZS, CA and SEE designed experiments and drew conclusions from the data. SEE supervised all experiments and analysis. JE, GT, EB, ED and YHC provided analysis pipelines for cyclic IF registration, segmentation and clustering steps. AA provided experimental and computational pipelines for single-cell ATAC sequencing. ZS, CA and SEE wrote the manuscript with input from all authors. SEE reviewed and edited the manuscript.

## Acknowledgements

We are grateful to the Biolibrary and the Histopathology Shared Resource (HSR) at OHSU for providing patient samples, specifically Aletha Letsch and Cheyenne Martin. The Biolibrary and HSR was supported by NIH grants P30 CA069533 and P30 CA069533 13S5 through the Knight Cancer Institute. We thank the Spellman lab and the Human Tumor Atlas Network (HTAN) for engaging discussions. We thank Stefanie Kaech Petrie and her team at the Advanced Light Microscopy Core at OHSU for their technical help. We thank Lindsey Cauthen for her feedback on the manuscript and for her revisions. We also thank Shelley Barton for providing feedback on the manuscript. This project was supported by funding (CEDAR3410918) from the Cancer Early Detection Advanced Research Centre at Oregon health & Science University, Knight Cancer Institute (S.E.E.).

